# Structural basis of sterol recognition by human hedgehog receptor PTCH1

**DOI:** 10.1101/508325

**Authors:** Chao Qi, Giulio Di Minin, Irene Vercellino, Anton Wutz, Volodymyr M. Korkhov

## Abstract

Hedgehog signaling is central in embryonic development and tissue regeneration. Disruption of the pathway is linked to genetic diseases and cancer. Binding of the secreted ligand, Sonic hedgehog (ShhN) to its receptor Patched (PTCH1) activates the signaling pathway. Here, we describe a 3.4 Å cryo-EM structure of the human PTCH1 bound to ShhN_C24II_, a modified hedgehog ligand mimicking its palmitoylated form. The membrane-embedded part of PTCH1 is surrounded by ten sterol molecules at the inner and outer lipid bilayer portion of the protein. The annular sterols interact at multiple sites with both the sterol sensing domain, SSD, and the SSD-like domain, SSDL, which are located on opposite sides of PTCH1. The structure reveals a possible route for sterol translocation across the lipid bilayer by PTCH1 and homologous transporters.

---

The hedgehog pathway is one of the key mechanisms of the developmental patterning in mammalian embryonic development, and is essential in tissue regeneration and homeostasis (1, 2). Disrupting mutations in the components of the hedgehog pathway are linked to diseases such as holoprosencephaly (3), Curry-Jones syndrome (4) and Gorlin syndrome (5). Furthermore, abnormal hedgehog pathway activation is associated with different forms of cancer, including medulloblastoma and basal cell carcinoma (6, 7).

The hedgehog morphogens are small secreted proteins. Three homologous hedgehog ligands have been identified in mammals: sonic (Shh), desert (Dhh) and indian (Ihh) hedgehog (8). Shh is the best studied of the three and is presumed to play the key role in the hedgehog pathway (9). The processed form of Shh is post-translationally palmitoylated at its N-terminal residue C24 (10). Furthermore, the protein undergoes autoproteolysis, leading to production of the mature ShhN fragment that contains a C-terminal cholesteryl moiety (11). The dually lipid-modified ShhN is released from the hedgehog-producing cells and can elicit short- and long-range hedgehog pathway activation in the developing embryo (12). The hedgehog pathway is activated in the hedgehog-receiving cells upon binding of the hedgehog ligand to its receptor, PTCH1, along with several recently identified co-receptors (13, 14). In the absence of ShhN, PTCH1 suppresses the activity of Smoothened (SMO), a G protein-coupled receptor (15). Upon binding of ShhN to PTCH1, SMO translocates to the Ellis-van Creveld (EvC) zone of the primary cilium (16), and activates GLI family of transcription factors, the hallmark of the hedgehog signaling pathway activation (17).

PTCH1 is a member of the Resistance-Nodulation-Division (RND) transporter family (18). It is a 12-transmembrane (TM) domain membrane protein, featuring two extended ectodomains that are involved in ligand recognition. The TM2-6 bundle of PTCH1 is annotated as the sterol-sensing domain (SSD) based on sequence similarity with similar domains in proteins involved in cholesterol homeostasis (18, 19). The SSD is presumed to be involved in the interaction with the hedgehog ligand, as suggested by recent structural and biochemical studies. Several recent reports described the interaction of ShhN with human PTCH1 in detail. Gong et al. obtained a high resolution cryo-EM structure of PTCH1 bound to a bacterially-expressed ShhN fragment, including residues 39-190 of human ShhN (20). The structure featured the ligand bound to the two ectodomains, with extreme N- and C-terminal portions of ShhN unresolved. Separately, Qi et al. solved a cryo-EM structure of PTCH1 bound to a native ShhN fragment that includes the palmitoylated N-terminus, including residues 24-188 (21). Contrary to the structure reported by Gong et al., this reconstruction showed the N-terminus of the ShhN ligand buried deep within the receptor, with palmitoyl moiety wedged within the “neck” region between the ectodomain and the membrane-embedded part of the receptor. Interestingly, the density of the hedgehog ligand in this reconstruction was inconsistent with that reported by Gong et al. This inconsistency was settled by the same authors who followed their study with a structure of a native ShhN-bound PTCH1 captured in a state where two PTCH1 molecules A and B were bound to ShhN via a ShhN-core interface (PCTH1A), and via the palmitoylated N-terminus of ShhN (PTCH1B) (22). This remarkable structure explained the seemingly contradictory results obtained previously with the monomeric PTCH1 molecules trapping the bound ligand in opposite orientations. Importantly, all three of the published structures of PTCH1 featured two extra densities corresponding to sterols bound at the lipid-protein interface of PTCH1 (20–22). Recently, a ligand-free PTCH1 structure was reported, featuring a dimeric arrangement of the mouse PTCH1 homologue. The ligand-free mouse PTCH1 displayed several lipid-like density elements along a duct connecting the outer leaflet SSD region and the lipid site in the ECD. Supported by biochemical evidence, the structure suggested a possible role of PTCH1 in cholesterol transport as a mechanism of SMO regulation (23).

Previous studies identified several ShhN variants that can be used for hedgehog pathway activation, with differing efficacies (24). The dually lipid-modified ShhN showed the highest potency of activation, whereas the unmodified ShhN fragment was least effective *in vitro*. Substitution of the C24 residue of the non-lipidated ShhN with two Ile residues (C24II-ShhN, here referred to as ShhN_C24II_) resulted in a substantial increase in the activity of the ShhN fragment. Since the discovery of this effect of the C24II substitution on ShhN, the ShhN_C24II_ has become an indispensable tool in a wide range of studies of the hedgehog pathway. Based on the close to native activity of the ShhN_C24II_ ligand, structural characterization of the PTCH1-ShhN_C24II_ complex should accurately describe the recognition of the native hedgehog ligand by its receptor, PTCH1.

To gain insights into the recognition of the modified ShhN_C24II_ hedgehog ligand by PTCH1, we solved the cryo-EM structure of the human PTCH1 in a complex with ShhN_C24II_. The protein construct PTCH1Δ combining the C-terminal truncation (residues 1-1188) and the Y645A mutation, devoid of the two PPXY motifs recognized by the HECT E3 ligases (25, 26), had excellent biochemical properties and was used for structure determination (Fig. 1A, Fig. S1C-E). The ligand, ShhN_C24II_, was generated by expression in *E.coli* as an N-terminally SUMO-tagged fusion (Fig. S1A-B). The purified ShhN_C24II_ protein was biologically active, with an apparent EC50 an order of magnitude lower than that of the recombinant human ShhN (Fig. 1B). The protein was mixed with the purified PTCH1Δ at a 2:1 molar ratio and subjected to cryo-EM analysis (Fig. S2, S3). The cryo-EM map obtained by image processing of the PTCH1Δ-ShhN_C24II_ dataset was refined to 3.4 Å resolution and was used for building a complete model of the complex (Fig. 1C-D, Fig. S3). The atomic model of PTCH1Δ-ShhN_C24II_ covers the sequence of PTCH1 residues D46-P1186, with an unresolved loop region comprising residues F614-T728. The model features six well resolved glycosylation sites, the extended N-terminus of the protein and a part of the intracellular loop 3 (IL3), which were missing in the previous reconstructions of PTCH1 (Fig. 3).

**Figure 1.**
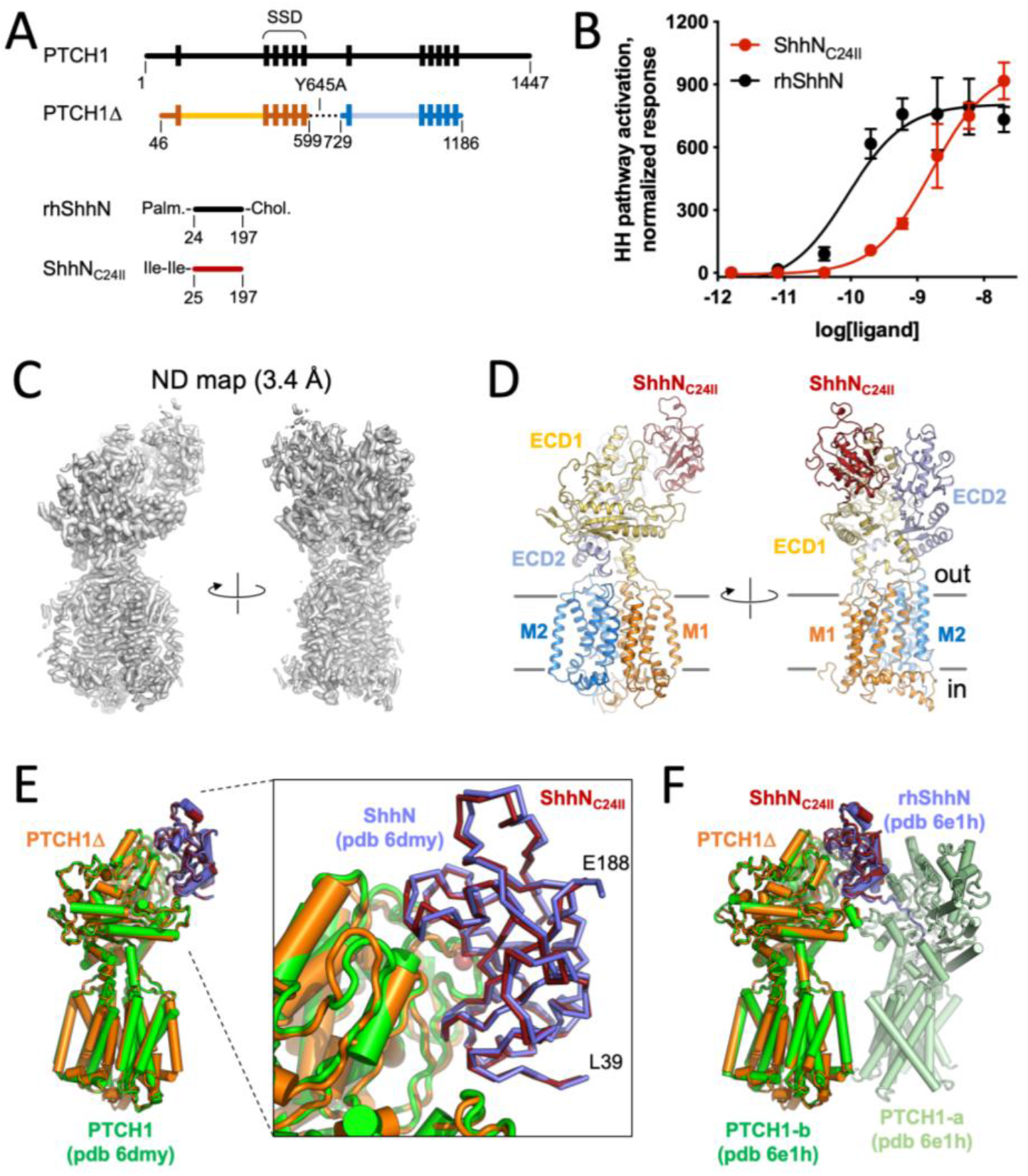
Structure of the PTCHlΔ-ShhN_C24II_ complex. A. A schematic representation of the constructs used in this study. For comparison, the wild-type full length proteins (PTCH1 and ShhN) are shown. B. The hedgehog pathway activation assay confirms the functionality of the ShhN_C24II_ preparation. NIH 3T3 cells were treated with the indicated concentrations of the ligands and hedgehog pathway activity was evaluated analyzing Gli1 mRNA levels by qPCR; error bars indicate S.D. (n=3). C. Density map of the reconstituted PTCHlΔ-ShhN_C24II_ complex. D. The views of the modelled complex corresponding to the views shown in (C). The key components of the complex are labelled, including the TM1-6 (“M1”), TM7-12 (“M2”) and the extracellular domains (ECD1-2). E-F. Comparison of the structure to previously solved PTCH1-ShhN (E) and PTCH1:rhShhN 2:1 complex (F). The bound ShhN_C24II_ perfectly aligns to the previously observed complexes mediated by the metal binding site interface of the ligand.

Comparisons of PTCH1Δ-ShhN_C24II_ and the two previously published reconstructions revealed that the unmodified ShhN binds in a near-identical manner (Fig. 1E-F). The N-terminus of ShhN_C24II_, similarly to that of ShhN, projects outside of the PTCH1 tunnel (Fig. 1E); the divalent metal binding site of the hedgehog ligand is close to the PTCH1 binding interface. Comparison to the rhShhN-bound PTCH1 structure shows that PTCH1Δ bound to the ShhNC24II matches the B-chain of the dimer (Fig. 1F).

Similar to the previously published reconstructions, our cryo-EM map contains four strong density elements likely corresponding to the bound sterol molecules (Fig. 2), here denoted as site 1 (at outer leaflet portion of the SSD; Fig. 2, Fig. 4A-B), site 2 (at the binding site formed by TM9, TM11 and TM12; Fig. 2, Fig. 4A-B), site 3 (buried within the ECD region; Fig. 2A, 2C) and site 4 (at the inner leaflet of the SSD; Fig. 2B-D). The “neck” region of the protein features a partial density element (site 0, Fig. 1A, inset), indicative of partial occupancy by an added sterol or a co-purified lipid-like molecule.

**Figure 2.**
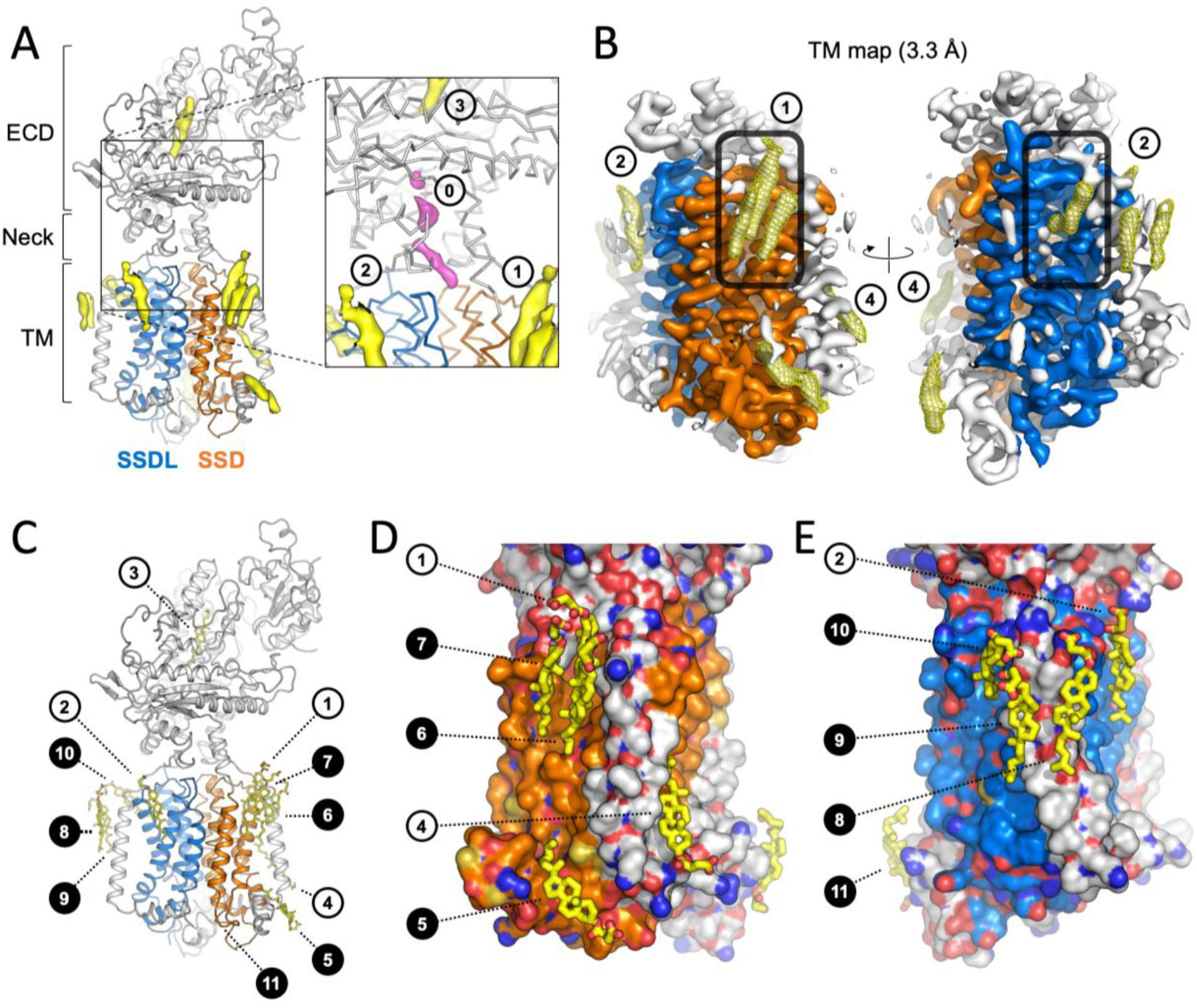
Bound sterol molecules revealed by the PTCHlΔ-ShhN_C24II_ structure. A. The 3D reconstruction features several well resolved density elements in the ECD region and surrounding the transmembrane domains of PTCH1Δ (isolated density are coloured yellow). The sterol sensing domain (SSD, TM2-6) and the SSD-like domain (SSDL, TM8-12) are coloured orange and blue. *Inset:* previously observed sterol sites are indicated as sites 1, 2 and 3; additional partial density is present in the “neck” region of PTCH1Δ (site 0, pink), suggesting partial occupancy of the site. B. Focused density map of the transmembrane region (TM map) indicates the well resolved sites 1, 2 and 4, as well as a number of novel sites, indicated with a mesh; the “pocket” region within the SSD and the corresponding are of SSDL is indicated by a rounded rectangle. C. The cholesteryl hemisuccinate (CHS) molecules were modelled into the well-defined density map regions (B), using 10σ cutoff of the unsharpened refined density map. The closed circles correspond to the novel sterol sites revealed by the PTCHlΔ-ShhN_C24II_ reconstruction. D-E. Views of the sterols surrounding the SSD and SSDL domains. Protein model is shown as surface, coloured according to atom type; carbon atoms in SSD and SSDL are coloured orange and blue, respectively.

The structure of PTCH1Δ-ShhN_C24II_ revealed a striking new feature: the presence of eight prominent density elements at the protein-lipid interface, additionally to sites 1, 2 and 4 (Fig. 2A-E, Fig. 4A-B). These novel positions of the bound sterols became apparent with the improvement of the density map corresponding to PTCH1 Δ. The focused refinement of the transmembrane domain of PTCH1Δ led to a density map at 3.3Å resolution, allowing us to describe the membrane-embedded portion of the protein with an unprecedented level of detail (Fig. 3B, Fig. S3; Movie S1). The positions of the membrane-embedded sterols surround the PTCH1Δ at sites within SSD (TM2-6), at the SSD-like region of the protein corresponding to TM8-12 (here referred to as “SSDL”), and at several sites not in direct contact with either SSD or SSDL (Fig. 2C-E).

**Figure 3.**
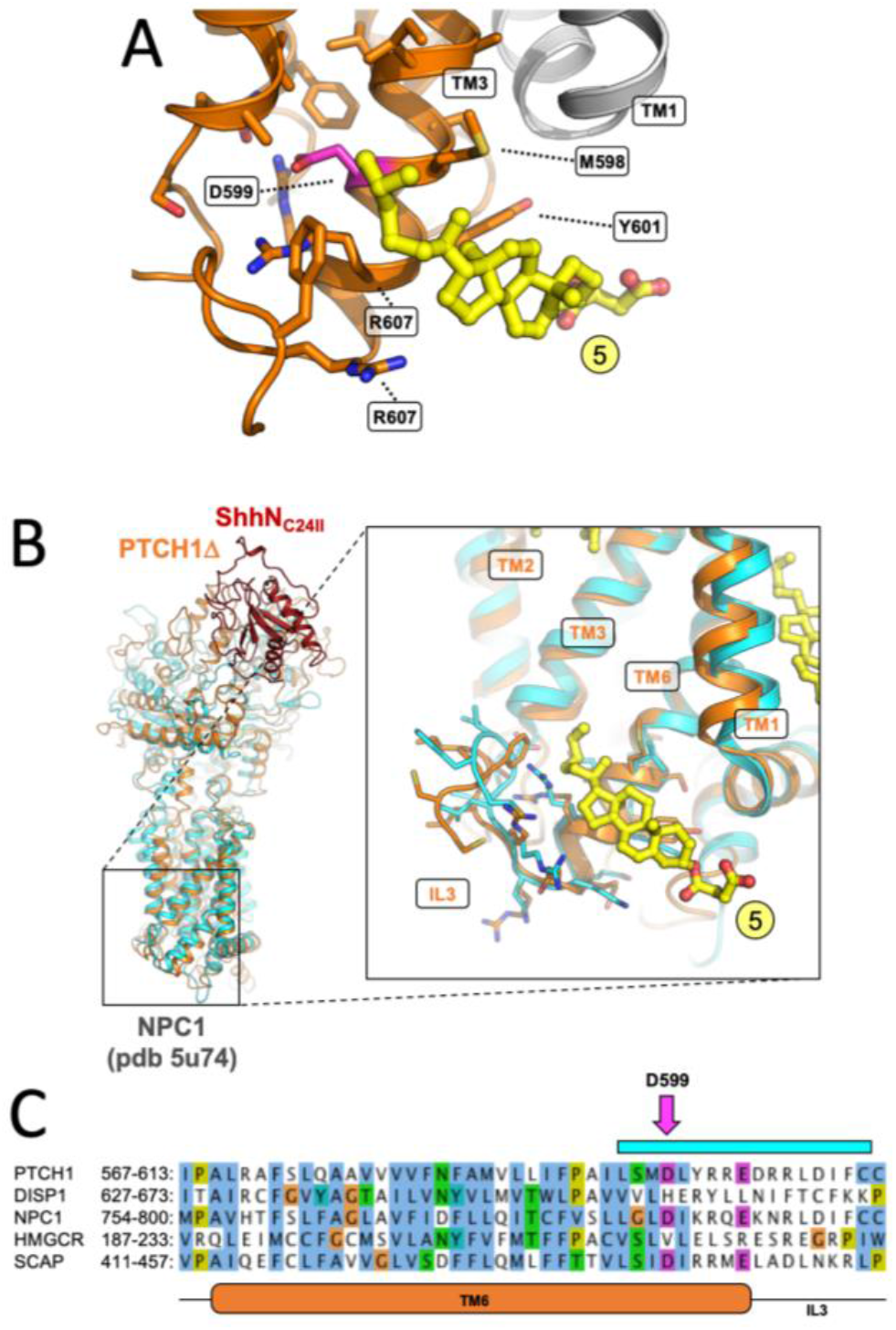
The inner leaflet SSD sterol site points at a functional role in PTCH1. A. The site 5 sterol (as defined in Fig. 2B; coloured yellow) is within contact of the residue D599, previously shown to be functionally critical in the SSDs of SCAP and Drosophila patched. B. A similar site, unoccupied by a sterol molecule, has been previously observed in the structure of cholesterol transporter NPC1, homologous to PTCH1. The PTCHlΔ-ShhN_C24II_ structure was aligned to the NPC1 X-ray structure (cyan). *Inset:* alignment of the two structures using the residues of TM6 shows a great degree of similarity in the region surrounding the site 5 sterol. C. Sequence alignment of key SSD-containing proteins, PTCH1, DISP1, NPC1, HMG CoA reductase and SCAP, shows the similarity between the elements involved in inner leaflet sterol binding site. The pink arrow indicates the residue D599 (shown in A). The cyan bar corresponds to residues shown with side-chains in (B). TM6 and intracellular loop 3 (IL3) of PTCH1 are indicated below the sequence alignment.

Sites 1, 6 and 7 occupy the outer leaflet “pocket” of the SSD (Fig. 2B-D). Site 5 is located at the inner leaflet portion of the SSD adjacent to the “pocket” (Fig. 2C-D, Fig. 3). Site 4 sterol at the SSD is sterically separated from the site 5 by the TM1 (Fig. 2C-D, Fig. 4A-B). The outer leaflet SSDL region is decorated with sterols at sites 2 and 10. Sites 8 and 9 are adjacent to the TM7. The site 11 sterol is located apart from other sterol molecules, bound to the N-terminal α-helix that precedes TM1 and lies parallel to the lipid bilayer plane (Fig. 2C, 2E).

**Figure 4.**
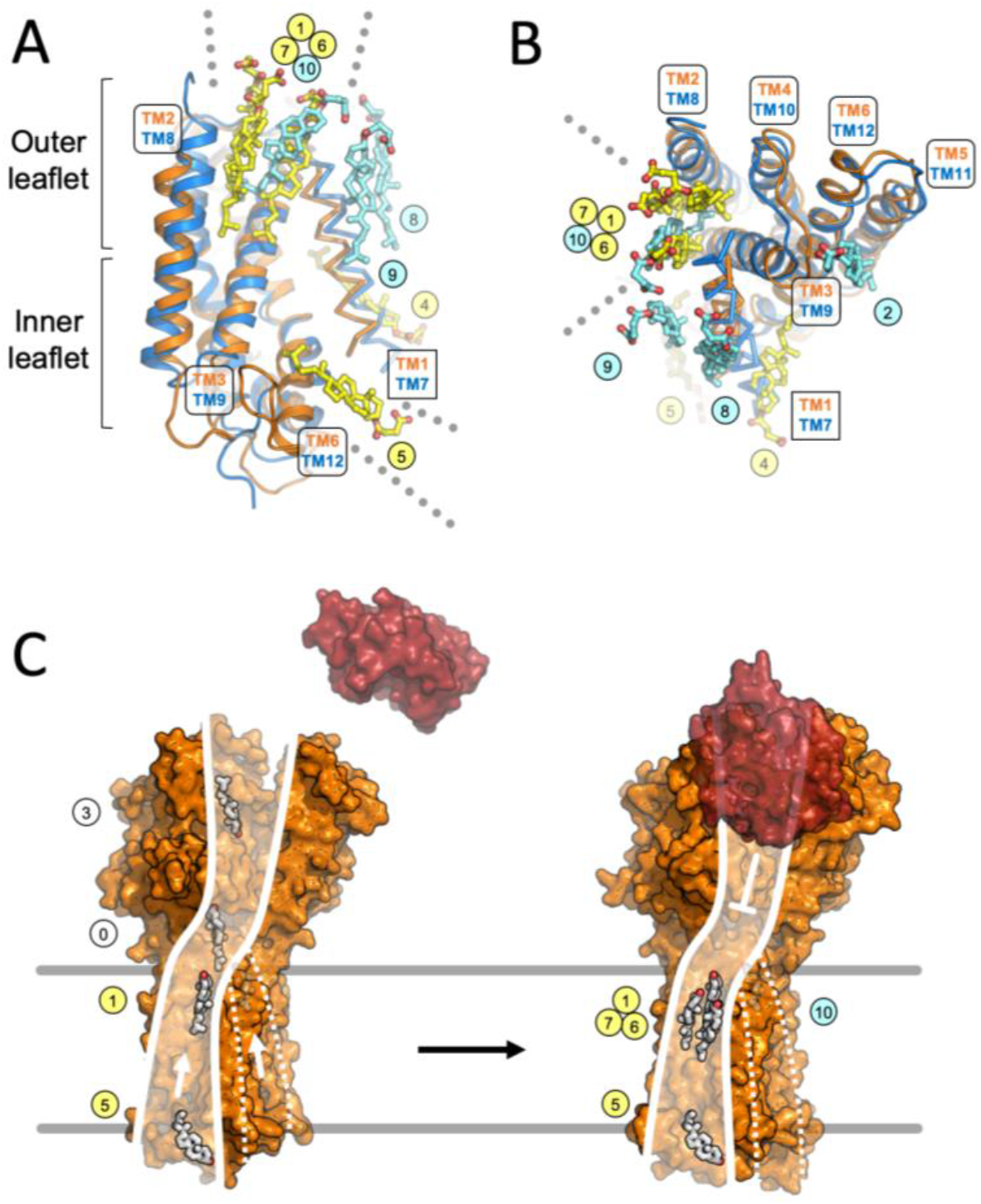
Sterol binding site overlap observed at the SSD and SSDL domains of PTCH1. A-B. The M1 and M2 regions of PTCH1Δ were aligned using the SSD (orange) and SSDL domain (blue). The SSD and SSDL regions are represented as cartoons; the transmembrane helices TM1 and TM7 are shown as ribbons. Sterols bound at the SSD (yellow) and SSDL region (light blue) are labelled using corresponding coloured circles. The outer leaflet site 1, 6 and 7 at the SSD and site 10 at the SSDL overlap. The dotted lines in A and B indicate access of sterols via the inner leaflet to site 5 (SSD) and via the outer leaflet to “pocket” sites 1, 6, 7 (SSD) and 10 (SSDL). C. Putative model of sterol translocation via PTCH1. *Left* – in the absence of hedgehog ligand (red); the protein (illustrated using PDB ID 6D4H) moves the inner leaflet sterols along the path formed by the SSD. *Right* – Binding of the ligand blocks this activity; the sterols accumulate at the SSD “pocket” region, evidenced by our 3D reconstruction. Presence of sterols at the SSDL, along with its similarity with SSD, may suggest a similar functional role (marked by the dashed lines).

The presence of the well resolved density of the sterol site 5 explains a previous observation of functional coupling between residues in this region and the activity of the SSD domain-containing proteins. It has been shown previously that D443N mutation in the SSD of SCAP, an ER-resident SSD-containing protein, disrupts the cholesterol sensitivity of the protein (27). A substitution of the equivalent residue D584 to an asparagine in *D. melanogaster* patched affected the ability of the protein to repress smoothened (28). Our reconstruction shows that the sterol bound at the site 5 is in immediate proximity to the D599 (Fig 2, Fig 3A), corresponding to D443 of the yeast SCAP or D584 of the drosophila patched. Comparison of PTCH1Δ to the X-ray structure of NPC1, a cholesterol transporter homologous to PTCH1, showed that a very similar structural motif is present in both proteins, likely forming a structurally conserved sterol binding site (Fig. 3B). Sequence alignment of PTCH1 and SSD domain containing proteins further confirmed the conservation of the residues in the region proximal to the site 5 sterol in PTCH1 and NPC1, two RND proteins presumed to be involved in cholesterol transport (Fig. 3C).

Structural alignment of the two pseudo-symmetric intramembrane halves of PTCH1Δ, using the SSD and SSDL regions for alignment, showed that most of the bound sterol molecules appear to be unique to the observed binding sites (Fig. 4A-B). However, there is a clear overlap between the “pocket” sterols: sites 1, 6 and 7 in the SSD and site 10 in the SSDL. This points to a possibility of a conserved role of both domains of the protein in interaction with the sterols, in line with the previously proposed model where cholesterol may access the inner tunnel at the “neck” region within PTCH1 from both sides (22). This would also be consistent with the presence of disease-linked mutations on both sides of PTCH1 (Fig. S5A). The mutations at the SSD side, more densely populated with bound sterol molecules, coincide with the residues directly involved in sterol binding (Fig. S5B). The mutations linked to holoprosencephaly-7, disrupting hedgehog signaling (3, 29), and the mutations linked to basal cell nevus syndrome, inducing aberrant hedgehog signaling (30), have been mapped to the SSD and the SSDL regions of PTCH1 (Fig. S5B-C), suggesting a likely functional conservation of the two domains.

In conclusion, the PTCH1 Δ-ShhN_C24II_ structure provides the missing clues for linking the sterol translocation pathway from the inner to the outer leaflet of the SSD (Fig. 4A-C). The existence of such a pathway was proposed by the recent reports (23), and our structure suggests the plausible routes for the sterol movement along the SSD surface from the inner to the outer leaflet of the lipid bilayer (Fig. 4C). The SSD domain of PTCH1 is capable of accumulating multiple sterol molecules in the “pocket”, as well as at the inner leaflet of the lipid bilayer. It is possible that the presence of ShhN_C24II_, a ligand that blocks the activity of PTCH1, establishes the conformation of the PTCH1 conducive to the accumulation of the sterols at the sites along the translocation pathway. The observed sterol sites are likely functionally coupled, as a number of mutations in this region abolish activity of the protein, as described above.

Furthermore, the presence of sterols at equivalent positions in the “pocket” region of the SSDL region of PTCH1 indicate that this domain may play a similar role to the SSD. Although the SSD regions have been defined based on sequence similarity to the SSD regions in HMG CoA reductase and SCAP (31), it has been noted previously that the distinction between the SSD and the regions corresponding to the SSDL domain in the RND family proteins may be somewhat arbitrary (18). The structural comparison of SSD and SSDL in PTCH1Δ confirms this suggestion and points at the functional similarity of these domains in sterol recognition (Fig. 4A-B).

Several well resolved sterol densities are present in our 3D reconstruction at sites distinct from the SSD and SSDL (sites 8, 9 and 11). The sterols bound at these sites may play an important structural role, stabilizing the membrane-embedded parts of the protein. These sites may also have a functional role, either directly participating in the substrate flux mediated by PTCH1, or by modulating its activity in various lipidic environments within or outside of the cilia membrane. The role of each lipid interaction will require careful investigation, and our structure provides the framework for the future in depth analysis of the protein-lipid interactions in PTCH1 and other SSD domain-containing membrane proteins.

## Acknowledgments

We thank the Electron Microscopy Facility at PSI, Villigen (Elisabeth Mueller-Gubler, Takashi Ishikawa) for support. We thank the BioEM Lab service facility at the University of Basel for the support in cryo-EM data collection. We also thank the Electron Microscopy Core Facility at EMBL Heidelberg for the support and expertise in high resolution cryo-EM data collection. This study has been supported by the Swiss National Science Foundation (SNF Professorship, 150665) and the ETH Grants (ETH-29 15-1) grants to VMK. C.Q. performed the experiments, analysed the data, wrote the manuscript; G.D. designed and performed the experiments, analysed the data, wrote the manuscript; I.V. designed and performed the experiments; A.W. analysed the data, wrote the manuscript; V.M.K. designed and performed the experiments, analysed the data, wrote the manuscript.

