## Supplementary Materials for "Structural basis of sterol recognition by human hedgehog receptor PTCH1"

**This PDF file includes:**

Materials and Methods
Figs. S1 to S5
Movie S1
Table S1

### Materials and Methods

#### PTCH1 expression and purification

The synthetic DNA fragment (Genewiz) encoding the human wild-type full-length PTCH1 protein (UniProt ID: Q13635) was cloned into pcDNA3.1 vector modified to contain a C-terminal GFP-10xHis tag fusion protein. PTCH1 was previously shown to be recognized by the HECT E3 ubiquitin ligases, via two PPXY motifs in the C-terminus (residues 1313-1316) and in the cytosolic loop (residues 642-645) (32, 33). Disruption of these motifs was shown to reduce PTCH1 degradation (33). Thus, we designed the PTCH1 expression construct to maximize the protein yield by removing the two PPXY motifs; the C-terminally truncated version of the protein, PTCH1-C (residues 1-1188), and the construct PTCH1Δ (comprising the residues 1-1188, with a mutation Y645A) were generated by PCR. The latter was subcloned into a modified pACMV plasmid (34), resulting in an expression cassette containing the PTCH1Δ construct fused to the C-terminal 3C-YFP-twinStrep tag. This was used to generate a stable tetracyclin-inducible HEK293-GnTi$^{-}$ cell line expressing PTCH1Δ.

Small-scale expression tests were performed using HEK293F cells. Expression plasmids encoding GFP-tagged wild-type full-length PTCH1 and derivative constructs were transfected into the HEK293F cells grown in 6-well plates using a PEI-based method in 6-wells. Cells were harvested 48 h later by centrifugation, resuspended in buffer A (50 mM Tris, pH 8.0, 200 mM NaCl, 10 % glycerol), sonicated and solubilized using 1% dodecyl maltoside (DDM). The lysates were centrifuged for 30 min at 25000 x g (Eppendorf) and supernatants were analysed by SDS PAGE and in-gel fluorescence using Amersham Imager 600 (GE: 29083461).

For large scale expression, the PTCH1Δ cell line cultures were expanded to 5L in 2L Erlenmeyer flasks at 37°C in Protein Expression Medium (PEM) supplemented with 5% Fetal Calf Serum (FCS), penicillin/streptomycin, Glutamax and non-essential amino acids until cell density of 2 x $10^6$ cells/ml was reached, and the expression of the protein was induced by adding 2 μg/ml tetracycline and 5 mM sodium butyrate. Cells were grown for additional 48 h at 37°C and were subsequently collected by centrifugation at 500 x g, frozen and stored at -80°C until the day of experiment.

For protein purification, the frozen pellets of 5L of HEK293-GnTi$^{-}$ cells expressing PTCH1 were resuspended in 150 ml of buffer A supplemented with 1 mM PMSF and 1 tablet per 50 ml of Roche complete protease inhibitor cocktail (EDTA-free). Cells were homogenized using a Dounce homogenizer and were solubilized directly by addition of 1% DDM and 0.2% cholesteryl semisuccinate (CHS) at 4°C for 1h. The lysates were clarified by centrifugation (Ti45 rotor, 35000 rpm, 30 min). The supernatants were added to 3 ml of CNBr-sepharose resin coupled with a purified anti-GFP nanobody (35). Following 1 h incubation with slow rotation at 4°C, the resin was collected in a gravity column (BioRad), washed with 30 column volumes of buffer A containing DDM/CHS mixture (0.025% DDM / 0.005 % CHS) and with 10 column volumes of buffer A containing 0.1% digitonin. The protein was eluted from the resin by cleavage with 3C protease overnight at 4°C. The eluted fractions were passed through 1 ml of NiNTA resin to remove the 3C protease. The protein was further purified using size exclusion chromatography using Superose 6 Increase 10/300 GL column equilibrated in buffer B (50 mM Tris-HCl, pH 8.0, 100 mM NaCl, 0.1% digitonin). The eluted fractions corresponding to PTCH1 were concentrated to 5 mg/ml (~38 μM) using an Amicon Ultra-4 concentrator (100 kDa cut-off, Millipore), and used for cryo-EM grid freezing immediately.

### ShhN$_{C24II}$ expression and purification

The synthetic DNA fragment (Genewiz) encoding the human ShhN$_{C24II}$ protein (residues 25-197, modified at the N-terminus with a Cys24-Ile-Ile substitution; UniProt ID: Q15465), fused with the N-terminal 6xHis-SUMO tag, was cloned into pET28a plasmid. The plasmid was transformed into BL21(DE3) RIPL *E.coli* cells. The transformed cultures were grown in a shaking incubator at 37°C until OD600 of 0.8, at which point the temperature was switched to 30°C and the expression was induced with 1 mM IPTG. After 3 h cells were collected by centrifugation, frozen and stored at -80°C until the day of experiment.

ShhN$_{C24II}$-expressing *E.coli* pellets (0.5 L culture) were thawed, resuspended in 10 ml buffer C (50 mM Tris, pH 7.5, 200 mM NaCl) containing 25 mM imidazole, disrupted by sonication, and cleared using centrifugation with a bench-top Eppendorf centrifuge for 30 min at 25000 x g. The supernatant was added to 0.5 ml of NiNTA resin, incubated with rotation for 1 h. The resin was collected on a gravity column (Biorad), washed with 30 column volumes of buffer C containing 25 mM imidazole, followed by 10 column volumes of buffer C containing 50 mM imidazole. The protein was eluted with buffer C containing 250 mM imidazole. The eluted fractions were pooled, diluted 8-fold to reduce the concentration of imidazole and incubated with 250 µg of purified SUMO protease Ulp1 overnight at 4°C. The mixture was passed though 1 ml of immobilized NiNTA resin to remove uncleaved protein, excess cleaved SUMO tag and Ulp1, the flow-through was concentrated to 1 ml and applied to a Superdex 200 Increase 10/300 GL column. The fractions corresponding to ShhN$_{C24II}$ were pooled, supplemented with 10% glycerol, aliquoted into 100 µl batches, flash-frozen in liquid nitrogen and stored at -80°C. Prior to cryo-EM grid freezing the protein was thawed, desalted into buffer B and concentrated to a final concentration of ~76 µM using an Amicon Ultra-4 concentrator (10 kDa cut-off, Millipore).

### Hedgehog reporter assays

To evaluate hedgehog pathway activation qPCR assays were performed. Confluent cultures of NIH 3T3 cells were starved overnight in DMEM, and treated for 24 hours with different concentrations of either rhShhN or ShhN$_{C24II}$. Following incubation RNA was extracted using the RNeasy Mini Kit (Qiagen) and reverse transcribed using the QuantiTect Reverse Transcription Kit (Qiagen). Transcription of the mouse Gli1 gene was measured by qPCR using KAPA SYBR FAST (Sigma) on the LigthCycler 480 System (Roche). mRNA expression of the SDHA gene was used for normalization. The data (as sown in Fig. 1B) were presented as fold enrichment with respect to the untreated sample. The sequences for the gene-specific primers were as follows: Gli1 (FW: GAATTCGTGTGCCATTGGGG, RV: GGACTTCCGACAGCCTTCAA), SDHA (FW: TTCCGTGTGGGGAGTGTATTGC, RV: AGGTCTGTGTTCCAAACCATTCC). All qPCR experiments were performed in triplicate starting from independent cell cultures.

### Electron microscopy data acquisition

To prepare cryo-EM grids containing the PTCH1Δ-ShhN$_{C24II}$ complex, the purified components, PTCH1Δ and ShhN$_{C24II}$, were mixed at a 1:2 molar ratio, with a final concentration of PTCH1 of 2.5 mg/ml (20 µM). The complex was incubated for 10 min at room temperature, followed by 30 min incubation on ice. Aliquots of the protein mixture (3.5 µl) were applied to the glow-discharged

UltrAuFoil 1.2/1.3 Au 300-mesh grids. The grids were blotted for 3 s, plunge-frozen in liquid ethane using Vitrobot Mark IV (Thermo Fischer Scientific), and stored in liquid nitrogen until the day of high resolution data collection.

Cryo-EM data collection was performed using a Titan Krios electron microscope (Thermo Fischer Scientific) equipped with a K2 Summit direct electron detector (Gatan) and a GIF-Quantum energy filter (slit width of 20 eV) at EMBL Heidelberg. The micrographs were recorded in counting mode with pixel size of 0.814 Å/pixel using SerialEM. The defocus range was set from -0.6 to -2.5 μm. Each micrograph was dose-fractionated to 40 frames with a total exposure time of 8 s, resulting in a total dose of ~44.7 $e^-$/Å$^2$.

The cryo-EM dataset for PTCH1Δ-apo (ligand-free) was collected at BioEM lab, C-Cina (University of Basel), using a similar set up (Titan Krios, K2 Camera, Quantum GIF) and methodology. The pixel size was 0.831 Å/pixel, the defocus range was -0.8 to -3 μm; each micrograph was dose-fractionated to 40 frames with a total exposure time of 10 s, resulting in a total dose of 75 $e^-$/Å$^2$.

**Cryo-EM image analysis**

A total of 7628 cryo-EM movies were motion corrected using MotionCor2 (36). Contrast transfer function (CTF) was estimated using Gctf on non-dose-weighted aligned images (37); micrographs with estimated resolution exceeding 4 Å were discarded, leaving 3260 micrographs for downstream processing. The PTCH1Δ-$ShhN_{C24II}$ particles were autopicked in relion-2.1.0 using templates generated by 2D classification of 1306 manually picked particles (38), resulting in a selection of 707620 particles. After several rounds of 2D classification, 430811 particles were selected for 3D classification. The procedure for 3D classification with 4 classes was performed in relion-2.1.0, using a 3D model of PTCH1Δ-apo low-pass filtered to 30 Å. The initial model was generated *de novo* in cisTEM using a dataset composed of 165533 PTCH1Δ-apo particles selected after 2D classification in relion-2.0. The best 3D class of PTCH1Δ-$ShhN_{C24II}$ complex consisted of 200679 particles, showed clear secondary structure elements (α-helices) and featured a bound molecule of the hedgehog ligand. This class was selected for 3D refinement. Refinement of this class gave a map with a resolution of 3.6 Å (based on FSC at 0.143). Reprocessing of this class using relion-3.0, including motion correction, CTF refinement and Bayesian particle polishing (39) resulted in a new map with a resolution of 3.5 Å ("F"). Further refinement was performed using the mask that excluded the detergent micelle, continuing from the last iteration. This resulted in a map at 3.4 Å resolution. The same procedure was repeated substituting the mask with a detergent and ectodomain-free mask, resulting in a 3.3 Å resolution map after postprocessing. Postprocessing was performed in relion-3.0 using a B factor of -50 Å$^2$. The processing steps and map improvements are illustrated in Fig. S2-3.

**Model building**

Model building was performed in COOT (40), using the previously solved structures of PTCH1 (PDB ID: 6DMY) and ShhN (PDB ID: 3N1R) for guidance. Bound sterol molecules were modelled as CHS (PDB code: Y01). The manually built model was refined using phenix.real_space_refine (41) implemented in PHENIX (42). For model validation, the atom positions of the refined model were randomly displaced by a maximum of 0.5 Å using the PDB tools in PHENIX. The derived perturbed model was subjected to real space refinement against one of the refined half maps (half-map1). Map vs model FSC comparison was made for the model against the corresponding half-map1 used in the refinement job, and for the same model versus the half-map2 that was not used during refinement (43). The model geometry was validated using MolProbity (44). Figures featuring the models and density maps were prepared using PyMol (45) and UCSF Chimera (46).

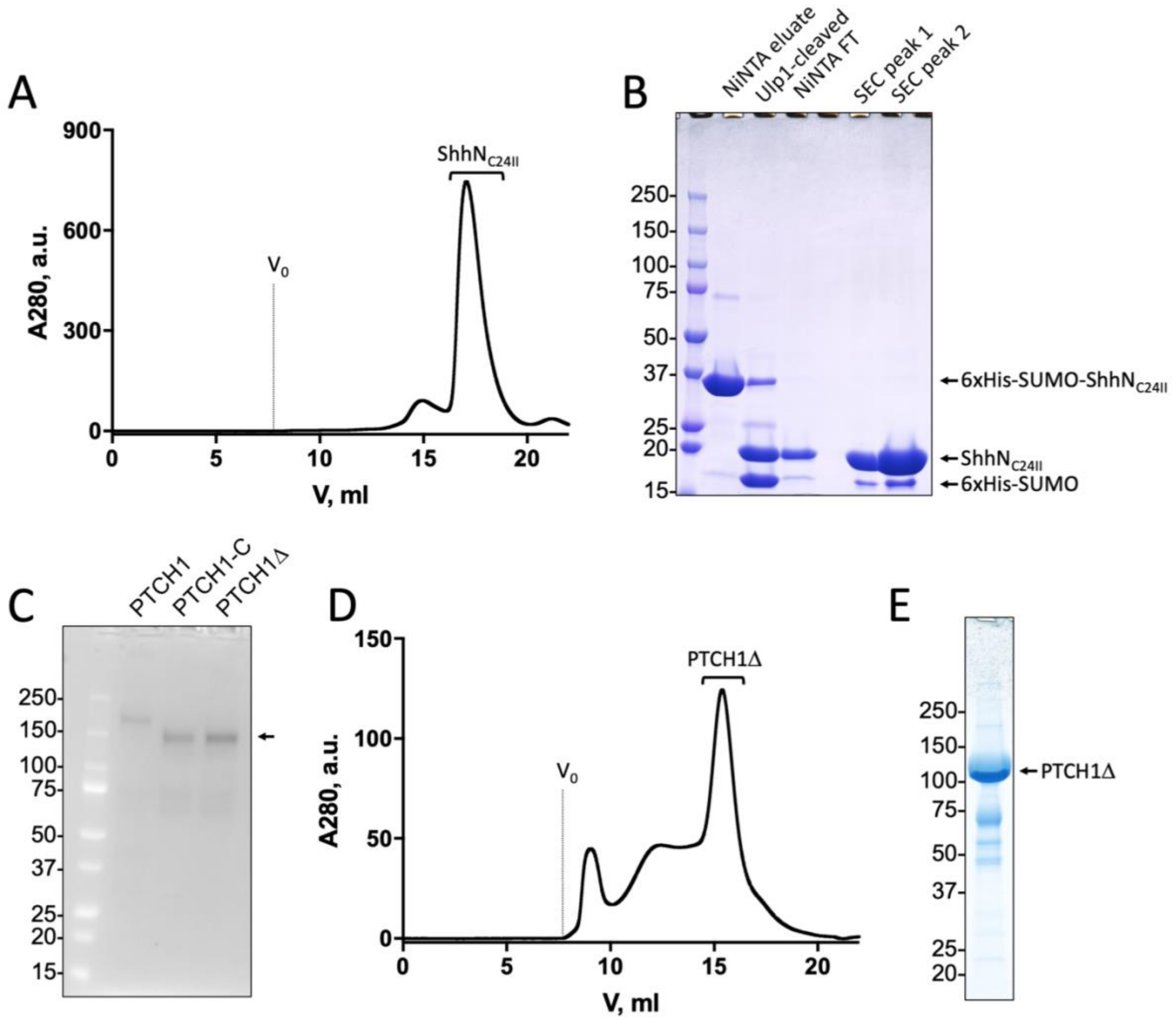


**Fig. S1. Purification of the hedgehog ligand and PTCH1.** A. Size exclusion chromatography (SEC) profile of the $ShhN_{C24II}$ protein; SEC was performed using Superdex 200 10/300 GL Increase column. The void volume is indicated by "$V_0$"; fractions chosen for complex reconstitution are indicated by a bracket. B. SDS PAGE analysis of the key stages of $ShhN_{C24II}$ purification. The fractions loaded on the gel were applied as indicated in the gel; Ulp1 is the SUMO protease; "NiNTA FT" is NiNTA flow-through after Ulp1 cleavage; "SEC Peak 1/2" indicate 3 μl and 10 μl of the same SEC peak (A), respectively. C. In gel fluorescence analysis of the SDS PAGE gel resolving the following samples: "PTCH1" – GFP-labeled full-length wild-type PTCH1; "PTCH1-C" – GFP-labeled C-terminal truncation of PTCH1 (residues 1-1188); "PTCH1Δ" the final construct used for structure determination (residues 1-1188, containing a Y645A mutation). Position of the truncated protein is indicated by an arrow. D. SEC profile of the purified PTCH1Δ preparation. SEC was performed using Superose 6 10/300 GL Increase column. E. SDS PAGE of the final PTCH1 sample.

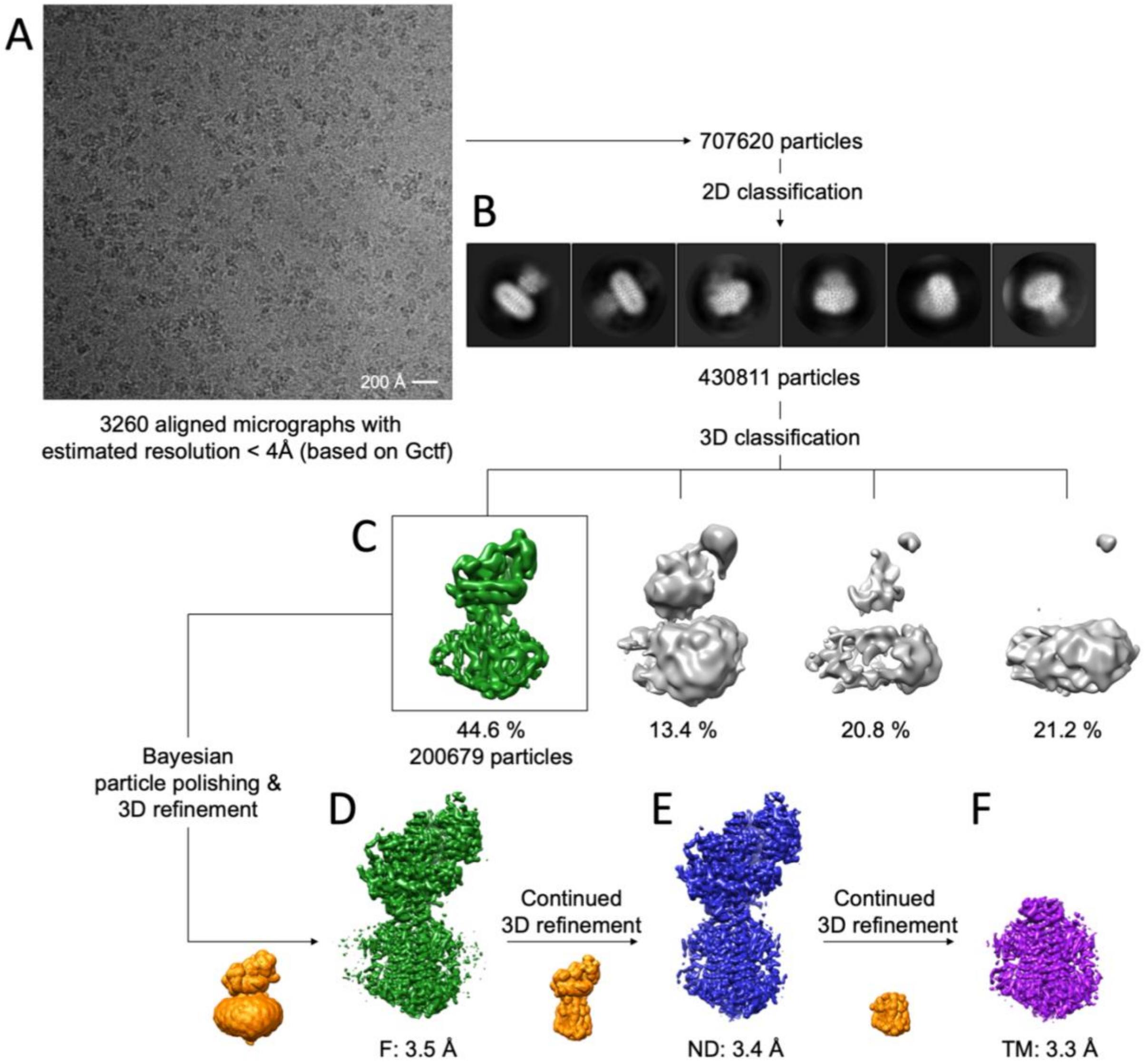


**Fig. S2. Cryo-EM and single particle analysis workflow.** A. The aligned averaged micrograph shows a high quality monodisperse sample of the PTCH1Δ-ShhN$_{C24II}$ complex in digitonin. B. The best 2D classes calculated in relion-2.0.b prior to the 3D classification step; 2D class features appear blurred. C. The 3D classification job with 4 classes resulted in a single class with clearly defined secondary structure elements. This class was subjected to further processing using relion-2.0.b and relion-3.0. D. Refinement of the best 3D class after Bayesian particle polishing in relion-3.0 (detailed in the "Material and Methods"), using a mask encompassing the complete complex (orange) resulted in a postprocessed density map at 3.5 Å resolution (full map, green, "F"; FSC 0.143). E. Continued refinement of the same dataset, substituting the full mask with a modified mask excluding the detergent micelle density resulted in a postprocessed map 3.4 Å resolution (no-detergent map, blue, "ND"). F. Continued refinement following the ND map, using the mask covering only the transmembrane region of PTCH1Δ resulted in the postprocessed map at 3.3 Å resolution (transmembrane map, magenta, "TM"). For each map, sharpening with a b-factor of -50 was used in the postprocessing step.

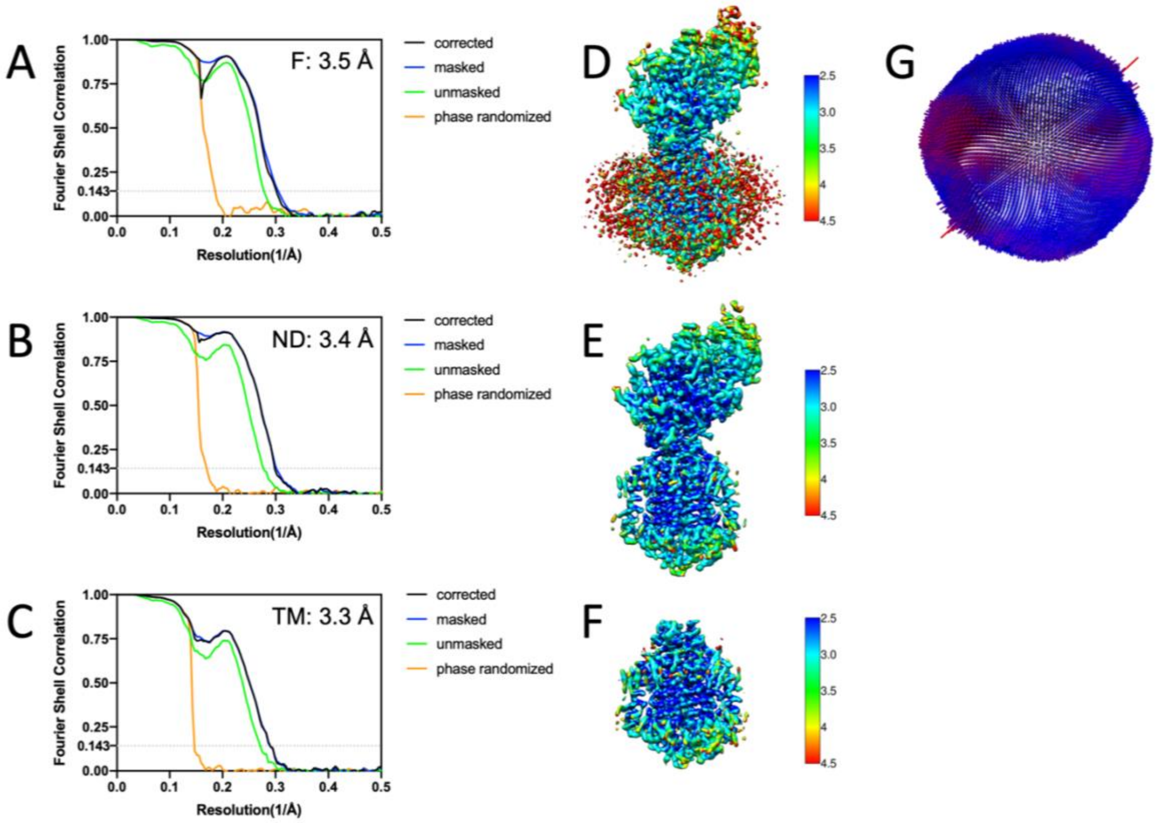


**Fig. S3. Fourier shell correlation (FSC) plots and 3D reconstruction details.** A-C. FSC plots for the three maps, "F", "ND" and "TM" (as defined in Fig. S2), calculated in relion-3.0. The dotted line indicates the 0.143 threshold ("gold standard FSC"). D-F. Local resolution estimates, calculated using ResMap implemented in relion-3.0. The scale bars indicate the resolution, scaled between 2.5 Å (blue) and 4.5 Å (red). G. Angular distribution of the refined dataset (calculated in relion-3.0 using the data in A and D).

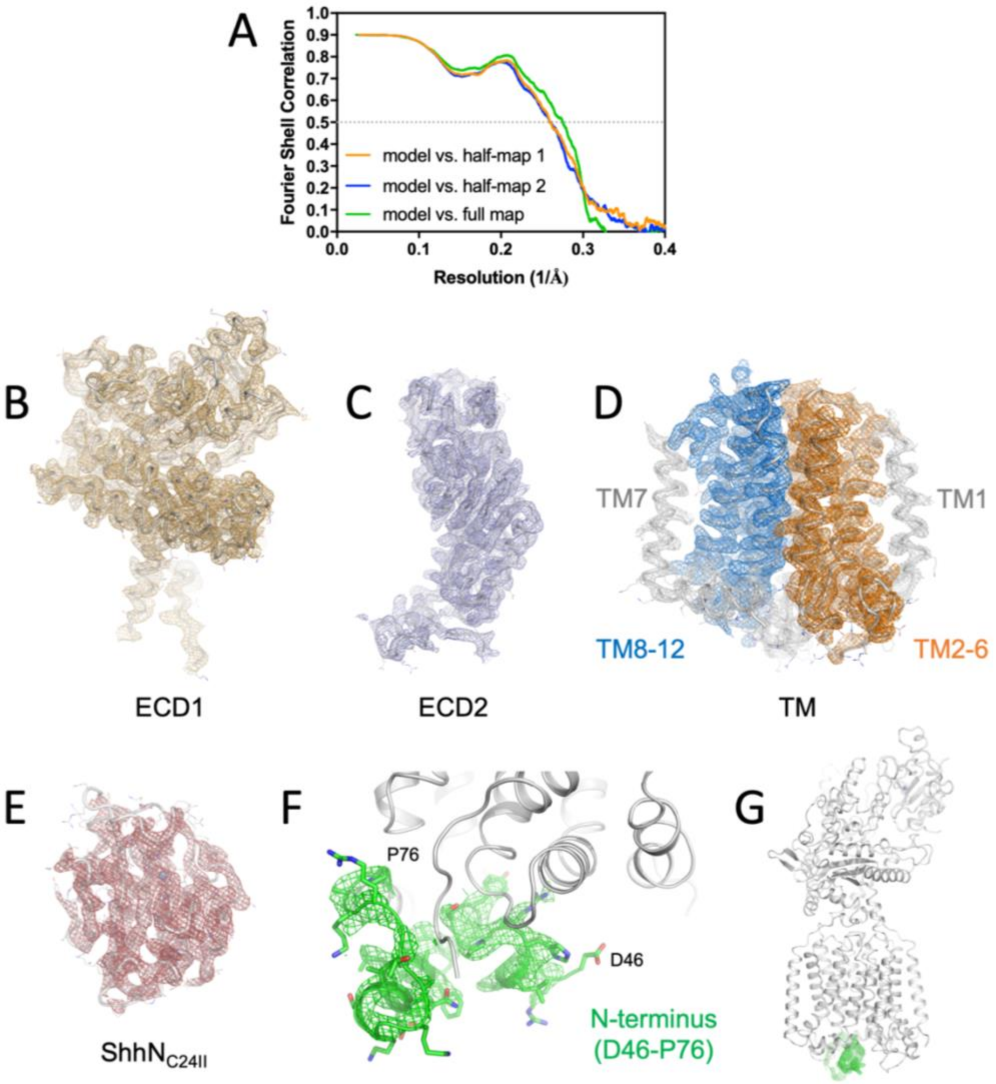


**Fig. S4. Features of the map and the atomic model of the protein.** A. Model to map FSC, calculated as detailed in "Materials and Methods"; the full statistics for the model are shown in Table S1. B-E. Individual parts of the map shown as mesh, including the regions corresponding to ectodomain 1 ("ECD1"), ectodomain 2 ("ECD2"), complete transmembrane domain bundle ("TM") and the modified hedgehog ligand ShhN$_{C24II}$. The protein model is shown as ribbon, with side-chain represented with lines (colored by atom type). F. The N-terminal residues of PTCH1Δ (D46-P76) resolved in our 3D reconstruction are shown as green mesh. G. The N-terminus forms an interface with the cytosolic side of the SSD and SSDL portions of the protein.

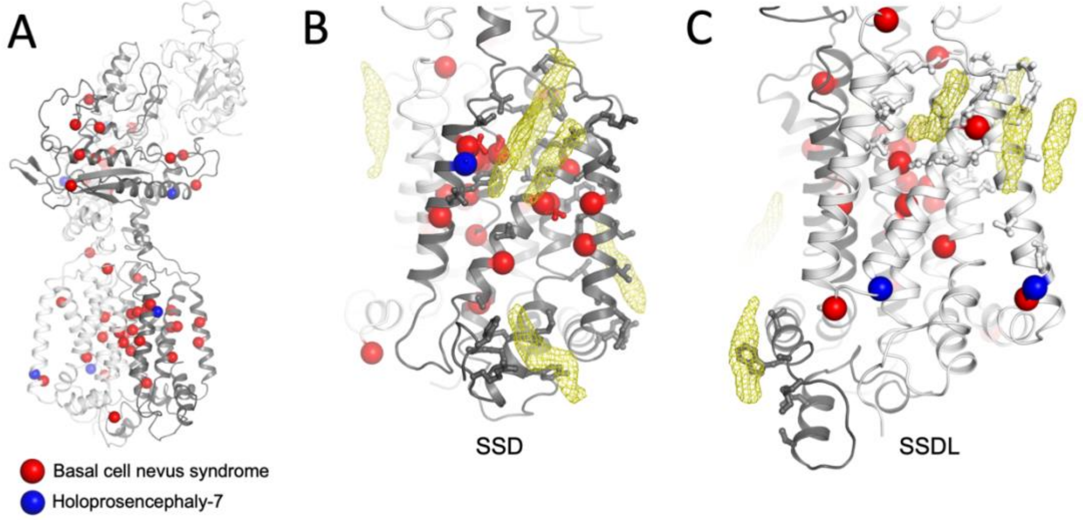


**Fig. S5. Disease linked mutations in PTCH1 are present within the SSD and SSDL regions of the protein.** A. Overview of the structure with the mutation sites linked to basal cell nevus syndrome (BCNS; red) and holoprosencephaly-7 (HPE-7; blue) indicated with spheres (C$\alpha$ atoms only). B-C. The views of the SSD (B) and SSDL (C) region show the positions of the disease-linked residues (spheres), with sterol binding site residue side-chains shown as sticks. Side chains that match residues linked to disease are coloured accordingly. Positions corresponding to the bound sterols in the TM density map are shown as yellow mesh.

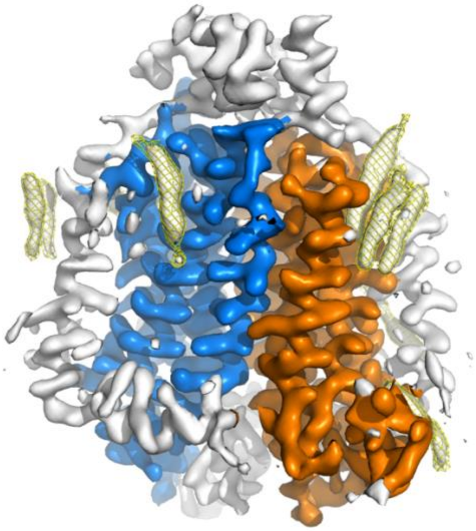

**Movie S1. The density elements corresponding to bound sterols in the PTCH1 TM focused map.** The movie was generated in Pymol (mset and util.mroll commands with 180 frames), using the TM map shown in Fig. 2B. The map is coloured according to Fig. 2B (SSD is coloured orange, SSDL is coloured blue). The positions of the sterol sites are indicated by yellow mesh over the density map.

**Table S1.** Cryo-EM data collection, single particle analysis and model building statistics.

| **Data collection** | |
|---|---|
| Instrument | FEI Titan Krios / Gatan K2 Summit / Quantum GIF |
| Magnification | 61425 (165kx) |
| Voltage (kV) | 300 |
| Electron exposure ($e^-/Å^2$) | 44.7 |
| Defocus range (um) | -0.8 to -2.4 |
| Pixel size (Å) | 0.814 |
| Resolution (Å; FSC 0.143) | 3.5 (F), 3.4 (ND), 3.3 (TM) |
| Number of particles | 200679 |
| **Model refinement** | |
| Model resolution (FSC 0.5; ND) | 3.7 |
| Map sharpening b-factor (Å) | -50 |
| Map CC | 0.771 |
| Model composition | |
| protein residues/ligands | 1195/19 |
| B factor ($Å^2$) | 102.52 |
| Bond length RMSD (Å) | 0.015 |
| Bond angle RMSD (°) | 1.434 |
| **Validation** | |
| MolProbity score | 1.82 |
| Clash score | 4.86 |
| Rotamer outliers (%) | 0.17 |
| Ramachandran plot | |
| Favored (%) | 89.49 |
| Allowed (%) | 10.31 |
| Disallowed (%) | 0.2 |
